# Antibiotic resistance response to sulfamethoxazole from the intracellular and extracellular DNA fractions of activated sludge

**DOI:** 10.1101/2022.11.14.516377

**Authors:** M. Martínez-Quintela, D. Calderón-Franco, M. C. M. van Loosdrecht, S. Suárez, F. Omil, D. G. Weissbrodt

## Abstract

In activated sludge, the antibiotic resistance genes (ARGs) can be present either in the intracellular (iDNA) or extracellular DNA fraction (exDNA). Recent advances in the exDNA extraction methodology allow a better profiling of the pool of ARGs. However, little is known about how stress conditions modify the distribution of ARGs between both DNA fractions. Here, we performed two batch tests for analyzing the effects of two different stress conditions, namely nutrient starvation and high concentrations of sulfamethoxazole (1, 10 and 150 mg L^-1^) in activated sludge. We tracked by qPCR the resulting relative abundances of four target genes, namely the universal 16S rRNA gene, the class 1 integron-integrase gene *intI1*, and the sulfonamide resistance genes *sul1* and *sul2* in both the iDNA and exDNA fractions. In the exDNA pool, unlike starvation, which provoked a decrease of 1-2 log_10_ [copies] ng DNA^-1^ in the concentration of *sul1* and *intI1*, the presence of sulfamethoxazole did not influence the abundances of *sul1* and *sul2*. However, high concentrations of sulfamethoxazole (150 mg L^- 1^) selected for microorganisms harboring *sul1* and, more remarkably, *sul2* genes in their iDNA during their exponential growth phase. The abundances of *intI1* and *sul1* were positively correlated in the exDNA fraction (r>0.7), whereas no significant correlation (*p*<0.05) between the abundance of these two genes was found in the iDNA fraction of the sludge. High SMX concentrations influenced the abundance of ARGs in the iDNA; their abundance in the exDNA was influenced by nutrient limitations. Further studies should consider the profiling of exDNA fractions because of the relationship between ARGs and mobile genetic elements. Besides, the surveillance of antimicrobial resistance is encouraged in wastewater treatment plants facing high antibiotic concentrations.

**Highlights:**

- Starvation caused a decrease in the exDNA concentration of the activated sludge.
- A positive correlation in the abundances of *intI1* and *sul1* was found in the exDNA fraction.
- High concentrations of SMX selected for antibiotic resistant microorganisms.

## 1. Introduction

Antibiotic consumption has greatly increased since 2000 worldwide, especially for human health purposes (Browne et al., 2021; Klein et al., 2018). Among them, sulfamethoxazole (SMX) is one of the most consumed antibiotics both by humans and animals (Wang et al., 2020). SMX inhibits the folic acid biosynthesis, hindering the bacterial growth (Jia et al., 2017). Its presence in wastewater is very well documented (Tran et al., 2018), and it has been recently included in the last update of the EU Watch list (EU2020/1161) as a concerning emerging contaminant. The overuse of antibiotics is considered the main driver of antibiotic resistance (AR), one of the major threats to human health in the current century.

The problem is not restricted to clinical settings since bacteria can move freely between human, animal, and natural environments. These compartments must be comprehensively taken into account when evaluating AR, following the One-Health approach (Miłobedzka et al., 2022). Wastewater treatment plants (WWTPs) should act as barriers to limit the spread of both, antibiotic resistant bacteria (ARB) and antibiotic resistance genes (ARGs), to the environment (Bürgmann et al., 2018; Calderón-Franco et al., 2022; Pallares-Vega et al., 2019). WWTP effluents, and wastewater catchment areas, are currently considered as one of the major release points of ARB and ARGs to the environment (Rizzo et al., 2013). Microorganisms present in WWTPs are considered as reservoirs of ARB since the mixture of animal and environmental bacteria coupled with the presence of trace concentrations of chemical stressors like antibiotics or heavy metals, creates a perfect environment for AR development and horizontal gene transfer (HGT) (Bengtsson-Palme et al., 2016; Karkman et al., 2018; Manaia et al., 2018). Although some studies have confirmed the capability of WWTPs to decrease the concentration of ARB and ARGs between influent and effluent (Lira et al., 2020; Pallares-Vega et al., 2021, 2019; Quintela-Baluja et al., 2019), others have reported a higher number of copies of some ARGs in WWTP effluents (Di Cesare et al., 2016; Rodriguez-Mozaz et al., 2015). Hultman et al. (2018) have detected some bacterial families in the WWTP effluent harboring one ARG (*tetM*) that was absent from the influent, suggesting possible HGT events in the activated sludge. The factors influencing the removal efficiency of ARB and ARGs, and HGT events are still unclear.

ARGs can be present either in the intracellular or extracellular DNA fractions. Extracellular DNA (exDNA) is commonly released from the intracellular DNA (iDNA) during cell lysis (passive release) or excreted from live cells (active release) (Calderón-Franco et al., 2020; Zarei-Baygi and Smith, 2021). Such exDNA can be present as a structural component in microbial aggregates, as part of the matrix of extracellular polymeric substances (EPS) (Costa et al., 2018; Rusanowska et al., 2019) or floating freely in the water body (Zhang et al., 2018). Recent advances in methods for the extraction of exDNA allow for a deeper analysis of the composition and distribution of both DNA fractions in WWTPs and natural environments (Calderón-Franco et al., 2021). The amount of exDNA detected in nutrient-rich wastewater environments is commonly of 1-2 orders of magnitude below iDNA (Calderón-Franco et al., 2021; Yuan et al., 2019). In contrast, a higher concentration (around 20%) of exDNA than iDNA has been detected in sediments in natural environments with nutrient scarcity (Mao et al., 2014). An increase (up to 2 log_10_ gene copies mL^-1^) of some ARGs (*ermB* and *sul1*) in the exDNA pool has been detected across a WWTP process, eventually discharged in the effluent (Calderón-Franco et al., 2022). The exDNA pool can therefore not be underestimated in AR surveillance in wastewater systems.

Once DNA is released from the bacterial cell, it is susceptible to be degraded abiotically or due to the action of extracellular nucleases (Torti et al., 2015). Several studies have stated that exDNA could persist without losing its integrity in natural environments for hours or days if it is free-floating, or even years if it is bounded to soils or surfaces (Nagler et al., 2018a; Nielsen et al., 2007; Thomas and Nielsen, 2005; Torti et al., 2015). Therefore, exDNA can also be considered a reservoir of ARGs and MGEs because of its high prevalence and stability.

Whether free or bound, exDNA can contribute to the spreading of ARGs via natural transformation in both pure and mixed cultures (Von Wintersdorff et al., 2016; Winter et al., 2021). Natural transformation is a mechanism of HGT in which bacterial cells in a competent state can take up, integrate, and functionally express genetic materials from foreign DNA (Blair et al., 2015; Winter et al., 2021). In the presence of stressors, this mechanism may be enhanced since some antibiotics and heavy metals can damage the cell membrane or DNA, creating the need to incorporate external DNA for reparation purposes (Zarei-Baygi and Smith, 2021). Antibiotics and heavy metals can also influence the ARG content in the iDNA fraction in high-density populations continuously exposed to them, such as those developing during biological treatments in WWTPs (Bengtsson-Palme et al., 2018). ARG transfer via HGT processes between colonies of *Escherichia coli* has been detected after applying 10 μg L^-1^ of tetracycline (Jutkina et al., 2016). Moreover, a list of antibiotics that may promote ARG selection processes in WWTPs in the range of 0.008 to 64 μg L^-1^ has been reported (Bengtsson-Palme and Larsson, 2016). The relationship between the distribution of the *sul1* and *sul2* genes, two of the most targeted ARGs related with sulfonamide resistance in WWTPs (Calderón-Franco et al., 2022; Di Cesare et al., 2016; Pallares-Vega et al., 2021, 2019), and the presence of SMX has not been previously assessed. The concentration of certain ARGs in both the iDNA and exDNA fractions can increase during the wastewater treatment process, suggesting HGT events through the WWTP (Calderón-Franco et al., 2022; Hultman et al., 2018). It is important to identify which WWTP conditions induce a shift in the ARB hosts and possible ARB reservoirs in order to delineate measures to reduce the dissemination of AR in the environment (Rice et al., 2020).

In this work, we evaluated the distribution of sulfonamide resistance genes (*sul1* and *sul2*), widely reported in wastewater, and the class 1 integron-integrase (*intI1*) genetic marker in both iDNA and exDNA fractions of activated sludge after a short-term exposure to sulfamethoxazole. Additionally, we aimed to elucidate resistance selection processes using the same target genes in activated sludge under different sulfamethoxazole concentrations.

## 2. Materials and Methods

### 2.1. Experimental setup

The inoculum for both experiments was obtained from the activated sludge of a WWTP located in Amsterdam West, The Netherlands. Each experiment was performed in 1 L Erlenmeyer flasks with 0.4 L of working volume at room temperature (~20 °C). Before inoculation, the biomass was filtered and washed to remove small particles. To ensure proper air transfer, flasks were placed in an incubator (Edmund Bühler) with a 120-130 rpm stirring rate. In each experiment, pH was manually controlled daily between 6.8 (initial) and 8.5 by addition of HCl at 1 mol L^-1^, to maintain the bacterial activity. A schematic representation of the experimental design is depicted in Figure 1.

**Figure 1.**
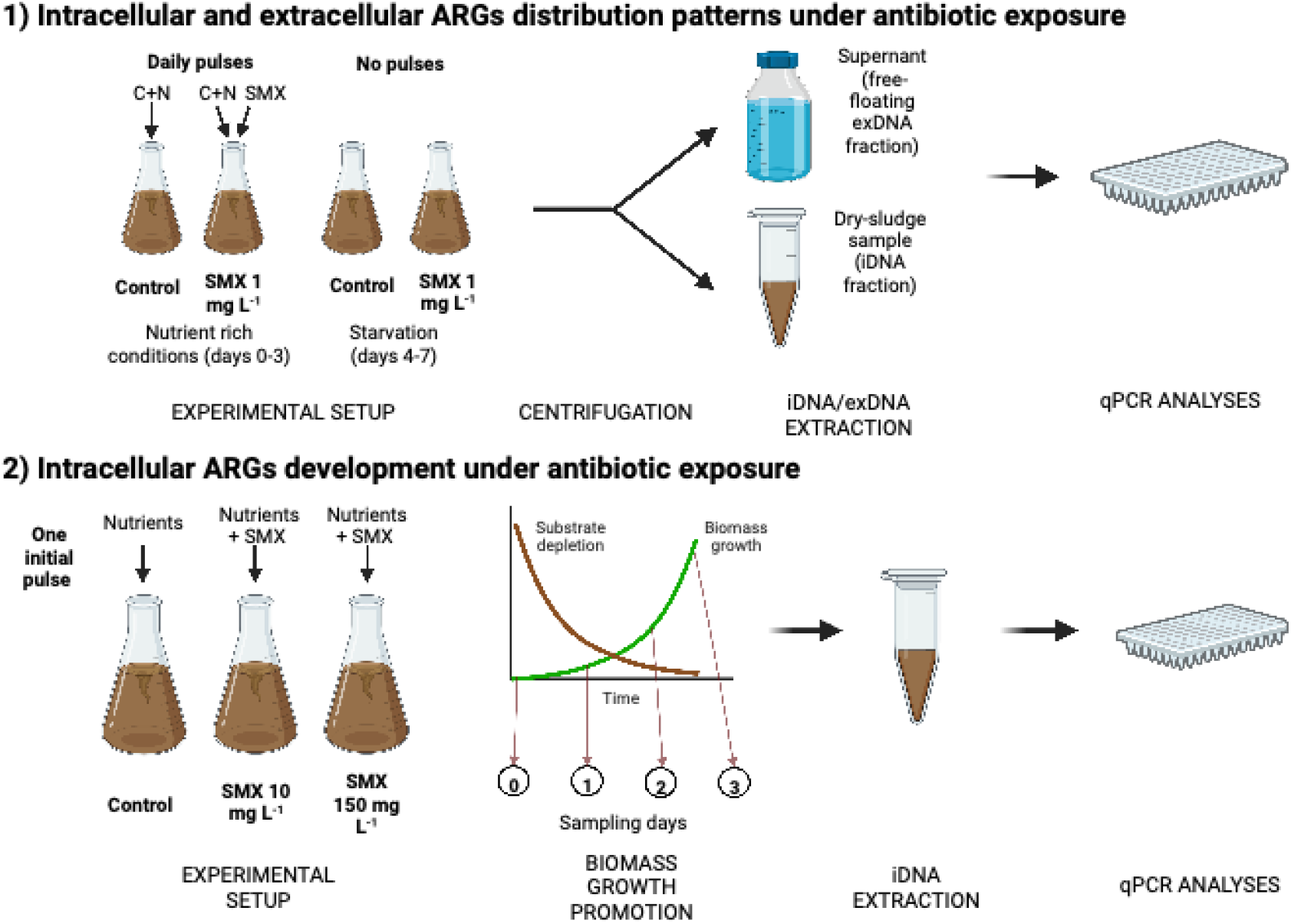
Schematic representation of both experimental designs. Created with BioRender.com

#### 2.1.1. Batch experiment 1: iDNA and exDNA distribution patterns under selective pressures

In this first experiment performed over one week, the effect of two different selective pressures (starvation and presence of SMX) on the distribution of ARGs in the iDNA and the exDNA fractions was analyzed.

##### Incubations and sampling points

Two different incubations were set up: one control (no antibiotic) and one supplied with SMX at an initial concentration of 1 mg L^-1^ in the mixed liquor. For each incubation, 3 sampling points were considered (1, 3, and 7 days) in duplicate, which implied the preparation of six flasks per incubation (total of 12 flasks). At each time point, the mixed liquor content of the whole flask (0.4 L) was used for performing the DNA extractions. Two additional flasks were prepared to measure the initial conditions (day 0) shared in both incubations.

##### Growth conditions and substrates pulses

In each flask, 0.4 L of activated sludge was inoculated. The initial concentration of volatile suspended solids (VSS) was about 4.40 g_VSS_ L^-1^. Flasks were continuously stirred through orbital shaking in an Edmund Bühler GmbH (K5-15) incubator to ensure a proper air transfer to maintain aerobic conditions. To maintain the biomass activity, sodium acetate (initial concentration in flask at 1.2 g COD L^-1^) and ammonium chloride (130 mg N-NH_4_^+^ L^- 1^) were pulsed to each flask daily until day 3 (included). Then, the supply of substrates to the microbial cultures was stopped to analyze the effect of nutrient starvation and cell lysis over the last 4 days of the experimental periods.

##### Antibiotic spikes

A stock solution of 7.8 g L^-1^ of SMX was prepared in methanol. 51 μL of this stock solution were spiked daily to obtain a SMX concentration of 1 mg L^-1^ until day 3 (included). A high concentration of antibiotic was used in comparison with those found in wastewater to enhance the antibiotic effect in the biomass. After day 3, antibiotic pulses were stopped to maintain starvation conditions.

##### Chemical and biomolecular analyses

Conventional parameters (chemical oxygen demand – COD, ammonium, nitrite) were analyzed at four different time points (0, 4, 8, and 24 h) during the first day, to check if the biomass was active. The concentration of total suspended solids (TSS) and VSS were analyzed on days 0 and 7 to measure biomass growth. The concentration of SMX was analyzed at the beginning of the experiment and before the addition of a new antibiotic pulse. The ARG content of both iDNA and exDNA fractions and the antibiotic concentration were analyzed at each time point (0, 1, 3 and 7 d) by qPCR and liquid chromatography, respectively, following the analytical methods described below.

#### 2.1.2. Batch experiment 2: Intracellular resistance genes development under antibiotic exposure

Considering the results of the first experiment, a second batch experiment was designed focusing only on the development of the intracellular ARG fraction under treatment with SMX over 3 days. The aim was to substantially promote the growth of the activated sludge, prior its exposure to SMX for a short time period, for monitoring the changes in the abundance of ARGs in the iDNA fraction.

##### Biomass acclimation

Initially, flasks were inoculated with 1.4 g_VSS_ L^-1^ of activated sludge biomass, which was about 4-times lower concentration than in the first experiment in order to monitor the effect of the antibiotic spiked during the exponential growth phase of the biomass. During the first week, the biomass was only fed with nutrients without antibiotic to acclimate it to a synthetic medium composed of easily biodegradable carbon, nitrogen, and phosphorus sources. To stimulate biomass growth, nutrients were provided in each flask in a single pulse containing a high concentration of sodium acetate (3 g COD L^-1^) and non-limiting concentrations of ammonium chloride (400 mg N-NH_4_^+^ L^-1^) and dihydrogen potassium phosphate (40 mg P-PO_4_^3-^ L^-1^).

##### Antibiotic supply

Once the nutrients were depleted, biomass was washed three times with tap water to renew the medium, and the antibiotic exposure was applied. For this purpose, 0.5 g_VSS_ L^-1^ of the pre-acclimatized biomass was used and fed with the same nutrient pulse as for the biomass cultivation period. A control (no antibiotic supply) and two different levels of antibiotic exposure were considered: 10 mg L^-1^ and 150 mg L^-1^. The spike was done only at the start of the experiment.

##### Chemical and biomolecular analyses

Conventional parameters (COD, ammonium and nitrite) and pH were followed at different times points (0, 24, 48, 72 h) during both the biomass cultivation and the antibiotic exposure periods. Biomass growth and SMX concentration were also checked daily during the first three days. Mixed liquor samples were collected each day to monitor the evolution of the targeted ARGs in the iDNA fraction.

### 2.2. iDNA and exDNA extraction

iDNA was extracted from 1-2 mL of mixed liquor from each flask. After centrifugation, a dry pellet of 0.10-0.25 g wet weight was obtained. The extraction of iDNA was performed using the NucleoSpin^®^ PowerSoil kit (Macherey-Nagel, USA) according to the manufacturer’s instructions. Each extraction of iDNA was normalized using 0.25 g of wet weight from each biomass pellet.

To retrieve exDNA, a sufficient volume (0.4 L) of mixed liquor was centrifugated and filtered through a PVDF 0.2 μm membrane filter. The sample volume for exDNA was first optimized to obtain a sufficient yield for performing qPCR analyses. The optimization results are described in section 3.1. The filtrate was loaded on a positively charged 1-mL diethylaminoethyl cellulose (DEAE-C) anion-exchange chromatographic column (BIA Separations, Slovenia). A liquid chromatography pump (Shimadzu, LC-8A) was used to load the sample into the column at a flow rate of 3 mL min^-1^, after equilibration of the column (procedure described in **Table 1**).

**Table 1.**
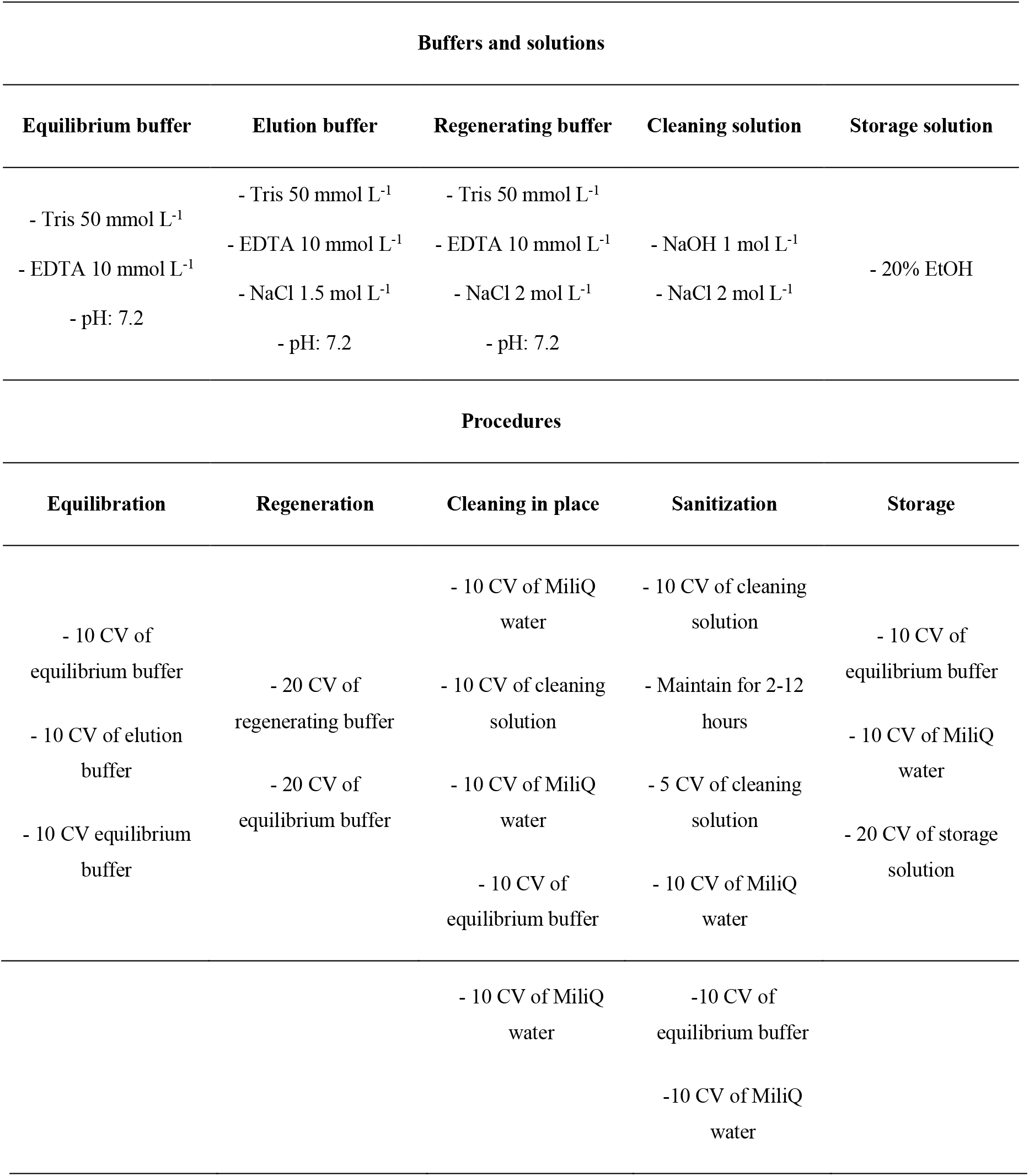
Compositions of the buffers and solutions, and procedures needed for column operation and maintenance for the isolation of exDNA. Buffers and solutions are composed by the mixture of the components listed. CV stands for column volume (1 mL).

After all the sample was loaded, the exDNA retained in the column was eluted during 20-30 min with an elution buffer (**Table 1**). Then, a volume of absolute ice-cold ethanol of 2-2.5 times the eluted volume was added to precipitate the DNA pellet. The supernatant was removed, and the pellet was washed two times with 70% ethanol, and the pellet was finally left to air dry. To improve the purity of the exDNA extract, the pellet was incubated with proteinase K for 2 h to remove residual proteins. The protein-digested pellet was finally purified using a GeneJET NGS Cleanup Kit (Thermo Scientific, USA).

Chromatographic columns require several procedures to maintain a proper operation. Buffers and procedures needed for column equilibration, regeneration, cleaning, and storage are described in **Table 1**. The equilibration procedure is required to prepare the column before the sample load. The regeneration procedure is performed after the elution of the sample. Cleaning and sanitization are washing procedures. The storage procedure is used to preserve the column integrity after the workout. This method has been previously described in Calderón-Franco et al. (2021).

### 2.3. Analytical methods

#### 2.3.1. Conventional parameters

Measurements of COD on filtered mixed liquor were performed using Hach Lange test kits LCK114 (Fisher Scientific; measurement range 150–1000 mg COD L^-1^) in the first batch experiment. During the second batch experiment, acetate concentrations were measured using a high-performance liquid chromatograph (HPLC) equipped with an Aminex HPX-87H column (BioRad, United States) maintained at 59°C and coupled to an ultraviolet detector at 210 nm (Waters United States). The eluent was phosphoric acid at 1.5 mmol L^−1^. Nitrogen compounds (ammonium and nitrite) were measured using a Gallery^TM^ Discrete Analyzer (ThermoFisher). Finally, TSS and VSS were measured according to Standard Methods (Rice et al., 2012).

#### 2.3.2. Analysis of sulfamethoxazole concentrations

The initial concentration of SMX and its evolution during the experiments was measured in an HPLC XLC-DAD (Jasco) equipped with a Gemini 3 μm C18 110A 150×4.6 mm column (Phenomenex). A binary solvent gradient consisting of 50 mmol L^−1^ of acetonitrile and phosphate (40:60, v/v) at pH 2.2 was used as the mobile phase. The injection volume was 100 μL, the column operated at 30°C, and the detector worked at 275 nm to detect SMX. The limit of quantification of the method was 50 μg SMX L^-1^. The method was previously described by González-Rodríguez et al. (2021).

#### 2.3.3. Quantitative polymerase chain reaction (qPCR)

Two of the most abundant sulfonamide resistance genes (*sul1* and *sul2*) described in wastewater were targeted. Additionally, the 16S rRNA gene was measured as bacterial reference and the class I integron-integrase gene *intI1* suspected of carrying some ARGs like *sul1* (Ghaly et al., 2021; Gillings et al., 2015) were analyzed.

All qPCRs were performed in a qTOWER3 Real-time PCR machine (Westburg, DE). qPCRs were conducted in 20 μL of reaction mixtures comprising: 10 μL of IQTM SYBR Green Supermix (Bio-Rad), 2 μL of DNA sample, 0.2 μL of forward primer, 0.2 μL of reverse primer (both at 10 μmol L^- 1^) and 7.6 μL of molecular grade water (Sigma Aldrich, USA). Three qPCR technical replicates were prepared for each sample. PCR conditions consisted of an initial denaturation at 95°C for 10 min followed by 40 cycles of DNA template denaturation for 15 s at 95°C, primer annealing for 30 s at 60°C for *intI1* and *sul1*, 55°C for 16S rRNA and 61°C for *sul2* genes, and primer extension at 72°C for 10 s. Forward and reverse primers are described in **Table 2.** A technical duplicate of at least 6 dilution series of synthetic DNA fragments (IDT, USA) (**Table S1**) containing the studied genes was included in each qPCR analysis to create the standard curve.

**Table 2.**
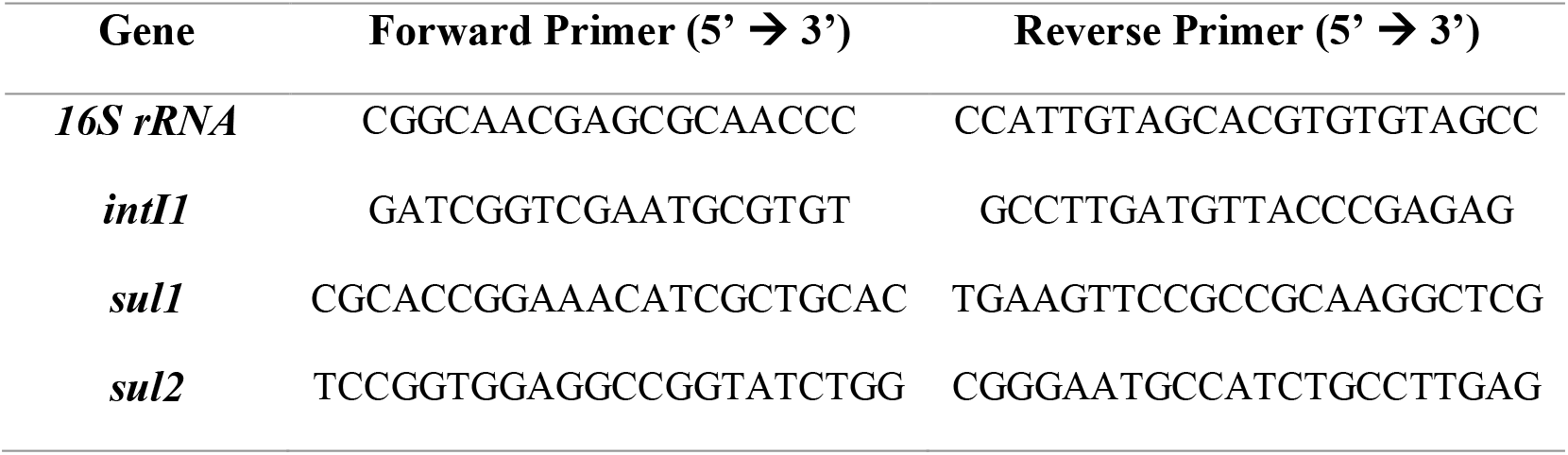
Primers used for qPCR analyses

### 2.4. Statistical analysis

To determine differences in the presence and abundance of ARGs in the microbiological samples, statistical tests were performed using the R software 4.1.0 and RStudio (https://www.rstudio.com/) at 95% confidence level (*p*<0.05). One-way ANOVA analyses were performed first to analyze significant differences among the samples. A post hoc Holm correction analysis was performed to adjust the *p-value* and determine which specific samples were significantly different. A Kruskal-Wallis test was applied when the normality assumption for applying the one-way ANOVA analysis was not satisfied.

## 3. Results and Discussion

### 3.1. Validation of the exDNA extraction method at different volumes and concentrations of mixed liquor

The exDNA extraction method was first tested to evaluate its sensibility to different sample volumes or biomass concentrations. Calderón-Franco et al. (2021) have shown that the anion-exchange chromatographic method using a diethylaminoethyl cellulose (DEAE-C) anion-exchange column can help recover more than 4 μg of exDNA from WWTP influent, effluent, and activated sludge using at least 1 L of sample. Here, a lower amount of sample volume was tested (0.25 and 0.5 L) as well as two different raw (~3.5 g_VSS_ L^-1^) and diluted (~1 g_VSS_ L^-1^) concentrations of activated sludge. Results on the retrieved exDNA concentrations and yields are depicted in **Table 3**.

**Table 3.** Concentrations and yields of exDNA extracted with different biomass concentrations and sample volumes (n.m. stands for not measured). Concentrations are expressed in ng of exDNA per μL of eluate and yields in ng of DNA.

|  | Initial volume<br>of activated<br>sludge | Raw sludge | Diluted sludge |
| --- | --- | --- | --- |
| exDNA<br>concentration<br>after extraction<br>(ng $\mu\text{L}^{-1}$) | 0.25 L | 100.00 | n.m. |
|  | 0.5 L | 104.00 | 45.20 |
|  | 1 L | 92.10 | 104.00 |
| exDNA yield<br>($\mu\text{g}$) | 0.25 L | 5.00 | n.m. |
|  | 0.5 L | 10.40 | 2.26 |
|  | 1 L | 18.42 | 5.20 |

As expected, the amount of exDNA obtained in each extraction decreased depending on the initial sample volume and biomass concentration. The amount of exDNA extracted was proportional to the amount of biomass in the sample: the yield (5.2 μg of DNA) obtained for 1 L of diluted sludge (*i.e*., from ~1 g_VSS_ of biomass) was similar to the one obtained for 0.25 L of raw sludge (5.0 μg of DNA) (*i.e*., from ~0.88 g_VSS_ of biomass). In all volumes and concentrations tested, a sufficient amount of exDNA was extracted. Such amounts (> 1 μg DNA) allowed further downstream analysis of the pool of exDNA by qPCR, and can also be used for metagenomics analysis of the resistome (not conducted here) (Calderón-Franco et al., 2021).

### 3.2. ARGs content was influenced by SMX in the iDNA and by starvation in the exDNA

The first batch experiment was run in flasks containing 4.4 g_VSS_ L^-1^ of activated sludge with pulse addition of acetate-based COD (at 1.2 g COD L^-1^) and ammonium (at 130 mg N-NH_4_^+^ L^-1^) as primary substrates. The consumptions of COD and NH_4_^+^ were followed on the first day of the experiment. Ammonium was consumed entirely after 1 d, but the soluble COD concentration remained at 200 mg L^-1^. Maybe the lack of phosphate or other trace mineral component limited the complete COD consumption. A volumetric maximum rate of COD consumption of 130 mg COD L^-1^ h^-1^ was measured in the first 8 h of the batch. Considering a theoretical COD:N:P ratio of 100:5:1 (m/m/m) for a balanced growth of ordinary heterotrophic organisms (OHOs) (Metcalf & Eddy, 2014), around 50 mg N-NH_4_^+^ would have been consumed per 1 g of COD_S_ of acetate. The conversion of ammonium could be due to both assimilation into OHO biomass and nitrification. The biomass concentration slightly increased from 4.4 g_VSS_ L^-1^ in the inoculum to ~4.9 g_VSS_ L^-1^ on day 7 in all experiments.

SMX was spiked at an initial concentration of 1 mg L^-1^ in the mixed liquor every day until day 3. Before the addition of a new pulse, the concentrations of SMX were always below the limit of quantification of the method (<50 μg L^-1^). According to the consumption of the conventional parameters (COD and N), the bacterial activity was not inhibited due to the antibiotic presence. The antibiotic measurements indicated a daily consumption of SMX, suggesting the biotransformation of the compound. Biodegradation rather than sorption of SMX has been previously reported (Alvarino et al., 2016). The biodegradation profile of SMX was also simulated with the Aquasim Software (Reichert, 1994) using kinetic and stoichiometric data (k_biol_SMX_= 3 L gVSS^-1^ d^-1^, K_s_ = 10 mg COD L^- 1^, Y_X/S_ = 0.5 g COD_X_ g^-1^ COD_S_, μ_max_= 5 d^-1^, b= 0.275 d^-1^) retrieved from literature for SMX biodegradation by OHO_S_ (Kennes-veiga et al., 2021) and from Activated Sludge Models (Henze et al., 1999). Simulations predicted a total consumption of SMX after 10 h of batch.

The amount of exDNA extracted varied significantly (p<0.05) throughout the experiment. Less exDNA was recovered on day 7 (3.95 ± 0.82 ng DNA mL^-1^) than on day 0 (13.53 ± 0.86 ng DNA mL^-1^). We hypothesize that the prolonged starvation conditions during the batch (where the water matrix was intentionally not renewed over the 7 days) led to the degradation of the residual exDNA or the starved biomass took up the exDNA as carbon source. DNA degradation can be caused by nucleases present in the sludge or released during cell lysis (Torti et al., 2015). In addition, bacterial inactivity could have resulted in lesser release of exDNA (Nagler et al., 2018b). Substantial analytical efforts will be needed in future for addressing the different mechanisms of exDNA release, uptake and degradation in microbial cultures.

Distributions of 16S rRNA, *sul1, sul2*, and *intI1* genes in both iDNA and exDNA fractions of the first set of batch experiments under control and SMX treatment conditions are depicted in **Figure 2**. Attending to the general trends in both incubations (control and SMX exposed flasks), very few differences in the abundance of the 16S rRNA gene were detected between days 0 and 7. This matched with the relatively low variation in biomass measured over the batch, which was inoculated at a relatively high concentration like an activated sludge tank. However, the rest of the targeted genes experienced a slight increase (~0.5 log_10_ [gene copies] ng DNA^-1^) in the number of copies in the iDNA fraction between these days.

**Figure 2.**
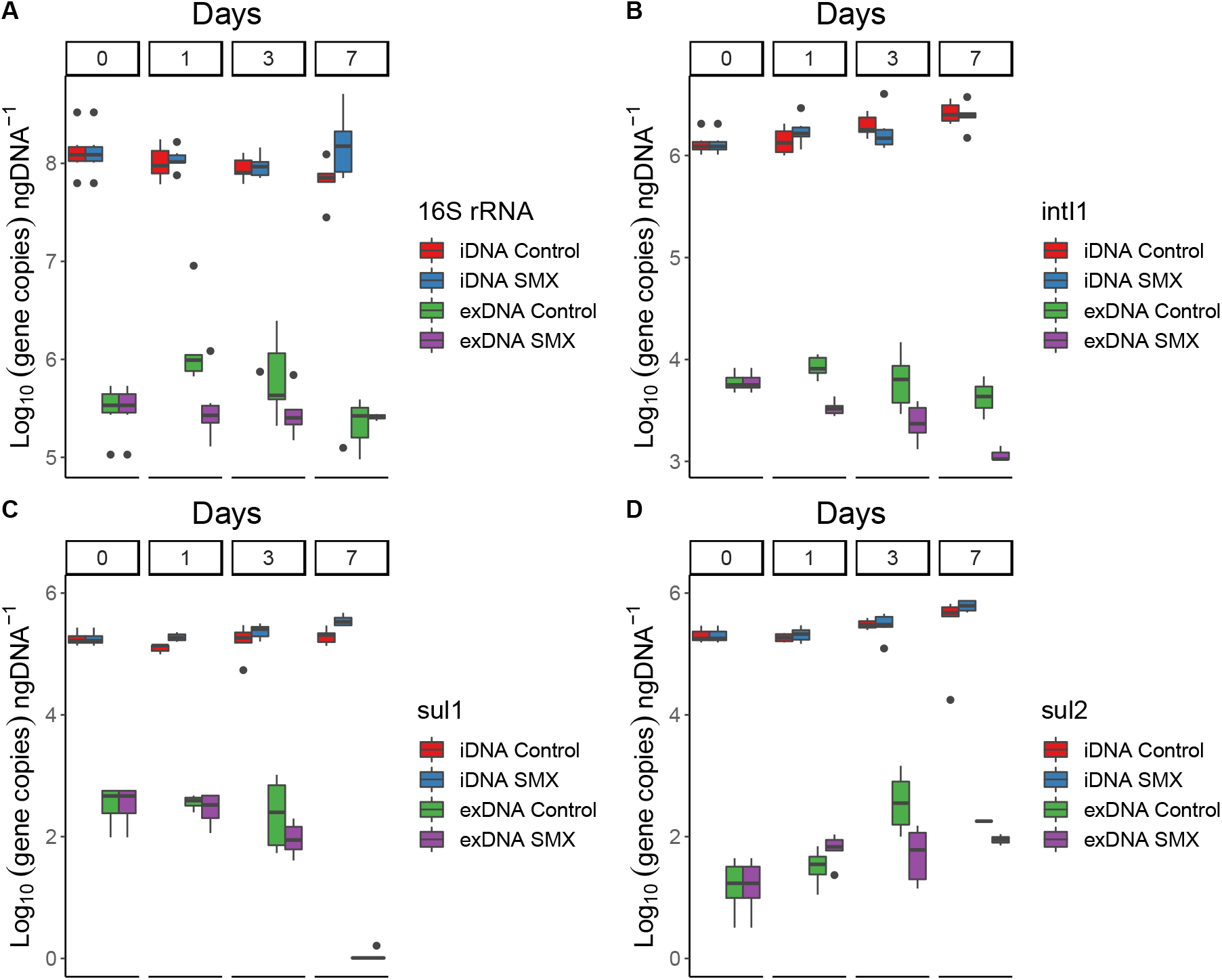
Distribution of gene copies of (a) 16S rRNA, (b) *intI1*, (c) *sul1*, and (d) *sul2* genes in both iDNA and exDNA fractions of the activated sludge over a week with and without (control) exposure to sulfamethoxazole (SMX). Nutrients and SMX addition were stopped after day 3. Results are normalized per ng of extracted DNA.

Focusing on the differences between the SMX exposed flasks and the control flasks, the number of copies of *sul1* and *sul2* in the iDNA fraction in the flasks exposed to SMX after 7 days was significantly higher (*p*<0.005) than on the starting day. This fact suggests a possible selection of microorganisms carrying those ARGs in their iDNA in the presence of the antibiotic. No significant differences (*p*<0.05) were found in the case of *intI1* between the different incubations.

In the exDNA fraction, the 16S rRNA gene barely varied from 5.5 log_10_ [gene copies] ng DNA^-1^ during these 7 days. There was a non-significant increase from 5.5 to 6 log_10_ [gene copies] ng DNA^-1^ on day 1 in the control flask, but the number of copies progressively dropped during the following days back to the initial value on day 7. The ANOVA did not show any significant difference (*p*<0.05) in the 16S rRNA gene copies in neither the control nor the SMX-exposed flasks.

The *intI1* and *sul1* genes exhibited similar profiles in the exDNA fraction of both the control and SMX treatment incubations: a progressive drop in their number of copies per ng of DNA was detected throughout the experimental period. A correlation between the number of copies per ng of DNA of each gene (*intI1* and *sul1*) were performed in both DNA fractions (**Fig. 3**). The gene concentrations correlated better in the exDNA fraction (r>0.7) than in the iDNA fraction (r<0.01), independent of the stress condition applied (either starvation or antibiotic exposure). It has been previously reported that class 1 integrons contain *sul1* in their gene cassettes (Ghaly et al., 2021). The correlation between the abundances of *intI1* and *sul1* genes on exDNA suggest that most of *sul1* genes present in this fraction were embedded in class 1 integrons. The experimental profiles highlighted significant differences among all days (*p*<0.005) for the *intI1* gene and *sul1* gene in the SMX-exposed flasks, especially between days 3 and 7. *sul1* was below the limit of detection on the last day. The number of copies of *sul2* in the exDNA fraction increased (>1 log_10_ ngDNA^-1^) significantly (*p*<0.005) from day 1 to day 7 to in the control flasks. In the SMX-exposed flasks, the significant (*p*<0.05) increase in *sul2* occurred between day 0 and 1, but in the following days the concentration was similar to the concentration of day 1 (~1.75 log_10_ gene copies ng DNA^-1^).

**Figure 3.**
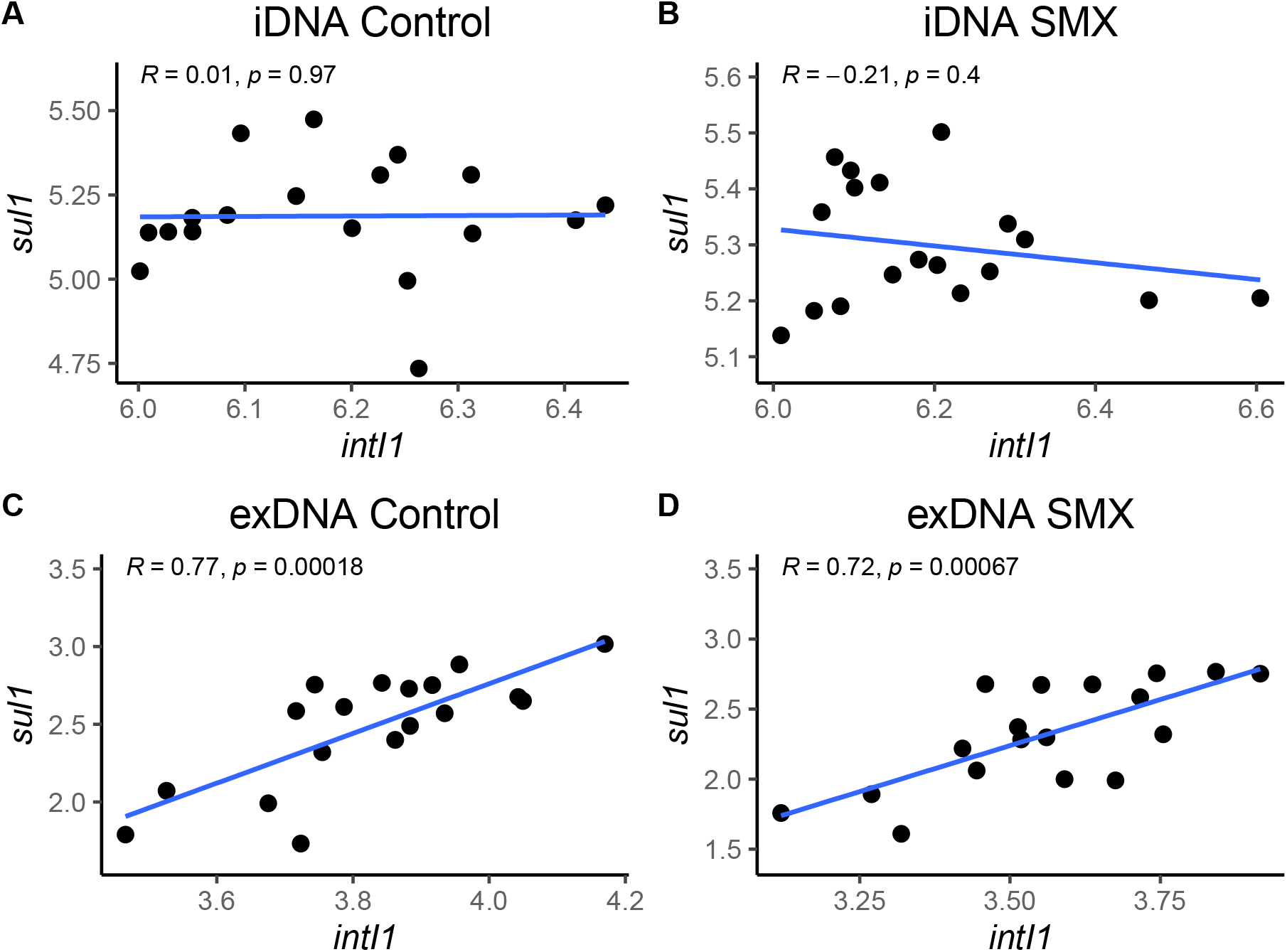
Correlations between the abundances of the *intI1* and *sul1* genes (given as log_10_ gene copies ng^- 1^ DNA) in the (a) iDNA fraction of control flasks, (b) iDNA fraction of the SMX-exposed flasks, (c) exDNA fraction of control flasks, and (d) exDNA fraction of SMX-exposed flasks.

The DNA release was not affected by the presence of the antibiotic but likely by the loss of bacterial activity. The exposure to 1 mg L^-1^ of sulfamethoxazole did neither significantly impact the bacterial activity nor the presence of *sul1* and *sul2* genes in exDNA. However, the presence of SMX selected for microorganisms carrying *sul1* and *sul2* in their iDNA over the non-resistant ones. As follow-up, we performed a second experiment to examine the ARGs concentration during the exponential phase of the biomass growth and focusing only on the iDNA fraction, since no significant differences were detected in the exDNA for the SMX resistance genes. The results are reported in the section 3.3 hereafter.

### 3.3. High SMX concentrations can select for antibiotic-resistant bacteria in activated sludge

The second batch experiment was inoculated at a 9-times lower concentration of previously acclimated biomass (0.5 g_VSS_ L^- 1^) for promoting growth and a selective enrichment during the experiment. Nutrients were supplied as one single initial pulse at a high concentration of acetate (3 g COD L^-1^) and non-limiting concentrations of ammonium (400 mg N-NH_4_^+^ L^-1^) and phosphate (40 mg P-PO_4_^3-^ L^-1^). The aim was to provide a sufficient amount of nutrients to promote the biomass growth over several days. Phosphate was also added here to avoid growth limitations. Nutrients were consumed progressively during the first two days. Acetate was set as the limiting compound (**Fig. 4a**). The high concentrations of SMX applied to the flasks (10 and 150 mg L^-1^), *i.e*., 10 to 150 times higher than in the previous experiment (1 mg L^-1^), did not affect the microbial activity. After 2 days, acetate was fully consumed, indicating that maybe the lack of phosphate was a limiting compound for the COD removal in the previous experiment. On day 2 growth and ammonium consumption also stopped (around 150 mg N-NH_4_^+^ L^-1^ were consumed in each flask). Likely, nitrifiers were more sensitive to the antibiotic than OHO_S_, as it was previously reported (Schmidt et al., 2012), since only traces of nitrite were detected (< 1 mg N-NO_2_^-^ L^-1^) in the SMX exposed flasks during the experiment. A slight increase in the ammonium concentration was detected between 48 h and 72 h (**Fig. 4b**), suggesting cell decay due to the lack of COD. Whereas Aquasim simulations predicted a full biodegradation of SMX over 30-50 h under the conditions of the second experiment (also considering an inhibition of growth by 50% under acute SMX treatment according to Katipoglu-Yazan et al. (2021)), the measured concentrations of SMX only slightly decreased during the first 3 days, namely 10-20% of the initial concentrations. In the flasks exposed to 10 mg L^-1^ of SMX, its concentration decreased down to 8 mg L^-1^ on day 3. In the case of flasks exposed to 150 mg L^-1^ of SMX, its final concentration after 3 days was around 140 mg L^-1^. During these experiments, the SMX concentrations remained high over the experimental growth periods of 2 days (**Fig. 4d**).

**Figure 4.**
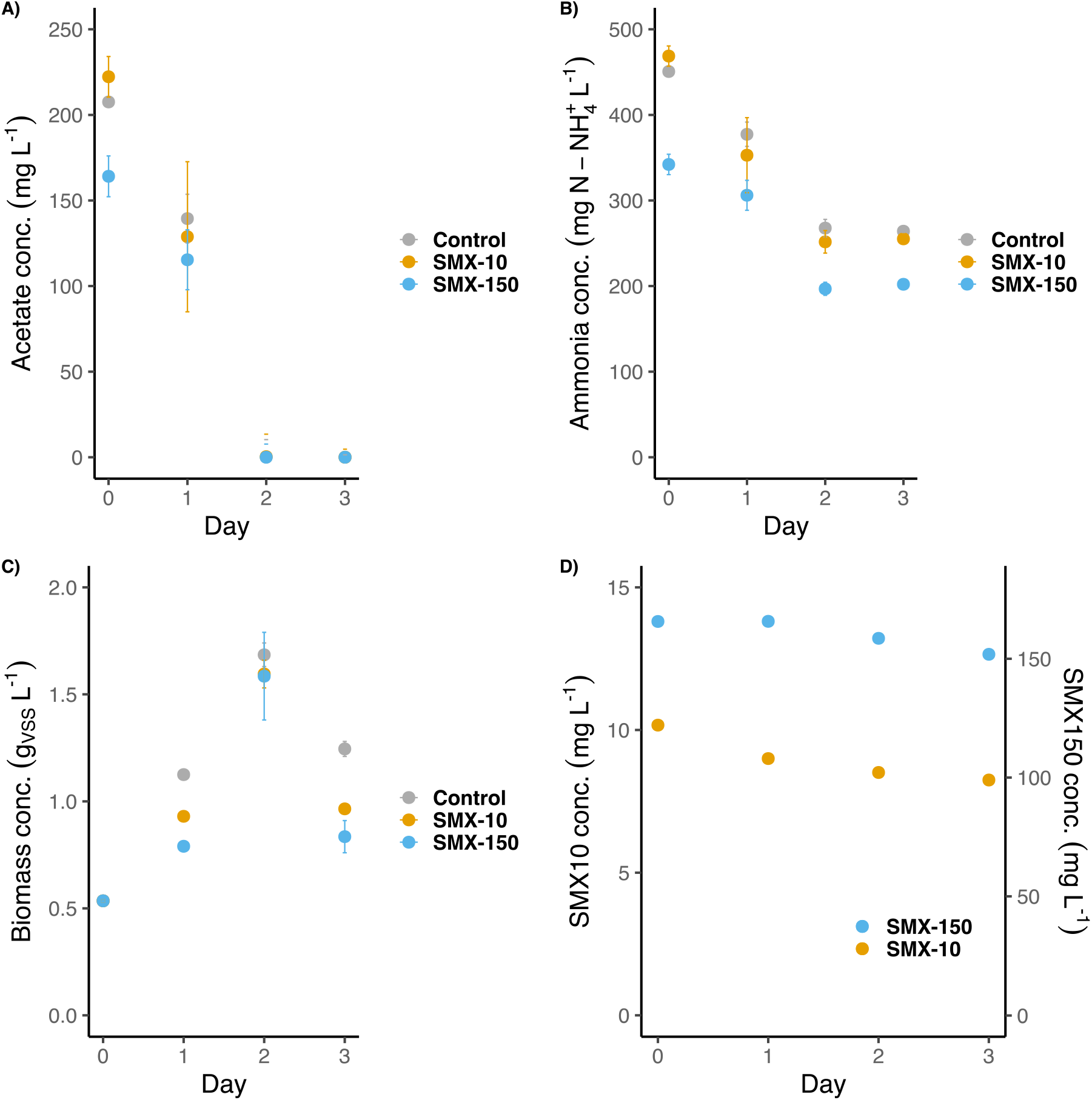
Evolution of the a) acetate concentration, b) ammonia concentration, c) biomass concentration and d) SMX concentration during the second batch experiment. SMX-10 and SMX-150 stand for the flasks exposed to 10 mg L^-1^ and 150 mg L^-1^ of SMX, respectively.

The biomass grew from its starting concentration (~0.5 g_VSS_ L^-1^) to a maximum of ~1.65 g_VSS_ L^-1^ on day 2 (**Fig. 4c**). Afterward, cell decay (and maybe some protozoa grazing) provoked a decrease in the biomass concentration to ~0.9 g_VSS_ L^-1^ on day 3. Although all flasks reached the same final biomass concentration, the growth rate patterns differed: flasks exposed to SMX grew slightly slower (0.011-0.016 g_VSS_ L^-1^ h^-1^) than the control one (0.025 g_VSS_ L^-1^ h^-1^). This suggested a partial inhibition of bacterial growth under antibiotic treatment. Still, the biomasses could quickly adapt to these antibiotic conditions and reach similar final concentrations as the non-exposed flasks.

According to the biomass dynamics, the qPCR analysis of the targeted genes was focused on the first 3 days (**Fig. 5)**. The abundance of the 16S rRNA gene followed in each flask a similar trend as the biomass concentration (**Fig. 5a**). Since the biomass of the control flask grew initially faster, the concentration of this gene on day 1 was higher than in the two SMX treatment flasks. However, on day 2, the 16S rRNA gene copies of the three flasks were similar (~7 log_10_ [copies] ng DNA^-1^), matching with the almost equal biomass concentrations obtained in the control and SMX exposed batches.

**Figure 5.**
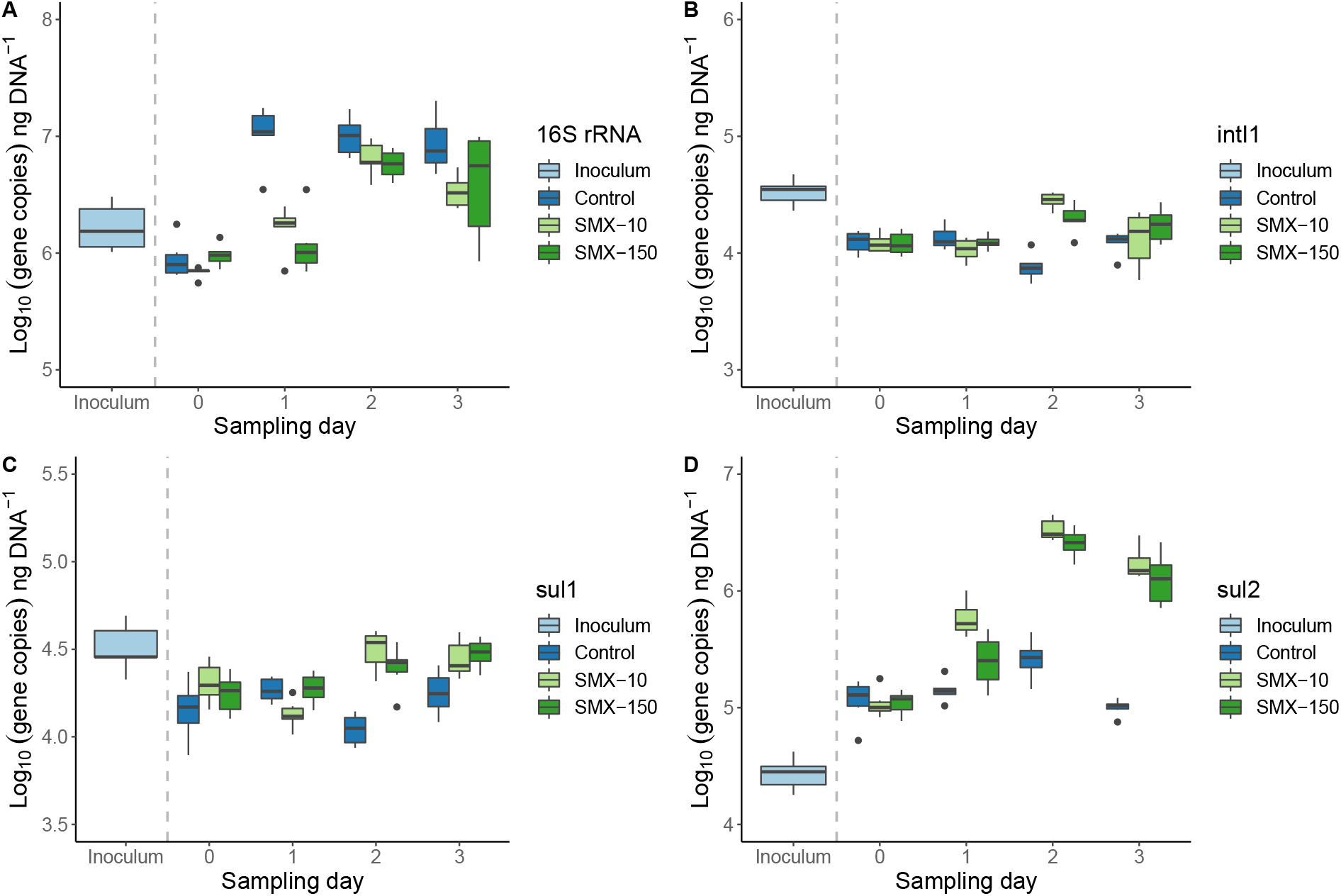
Distribution of gene copies of (a) 16S rRNA, (b) *intI1*, (c) *sul1*, and (d) *sul2* in the iDNA fraction of the activated sludge in the inoculum and the first 3 days of the experiment for three different concentrations of SMX: 0 (control), 10 and 150 mg L^-1^. Results are normalized per ng of extracted DNA.

The abundances of the *intI1* and *sul1* genes were very similar among the flasks (4.0-4.5 log_10_[copies] ng DNA^-1^) (**Fig. 5b-c**). However, they differed in their distribution over time. A slight significant (*p*<0.05) increment was detected on day 2 in *intI1* between the control and exposed flasks (from 4.0 to 4.5 log_10_[copies] ng DNA^-1^), but it was compensated on day 3. For *sul1*, the same significant (*p*<0.05) difference between the control and exposed flasks was detected on day 2. On day 3 the concentration of *sul1* in the control flasks decreased down to ~4.25 log_10_[copies] ng DNA^-1^. Unlike for the control and the SMX-10 flasks, the abundance of *sul1* significantly differed (*p*<0.005) between day 0 and day 3 in the SMX-150 flasks. The SMX treatments selected for microorganisms carrying *sul1* and *intI1* genes on their iDNA, *i.e*., potentially able to develop resistance and grow. Thus, the exposition to SMX resulted in an increased abundance of both *sul1* and *intI1* genes in the bacterial community.

The *sul2* concentration also differed depending on the presence of the antibiotic. In the control, the concentration barely varied from 5 log_10_[copies] ng DNA^-1^ (initial concentration at day 0) (**Fig. 5d**). In both flasks exposed to SMX, the abundance of *sul2* increased up to a value of 6.5 log_10_[copies] ng DNA^-1^ on day 2 (*p*<0.005). Such *sul2* concentration was maintained even along the cell decay that occurred between days 2 and 3. The high concentrations of SMX selected for bacteria that significantly carried *sul2*. The selection of microorganisms harboring *sul2* compared with those harboring *sul1* is remarkable. Since their resistance mechanism is the same (target protection) (Rizzo et al., 2013), a possible explanation for the large selection of *sul2* over *sul1* could be related to their different location. The *sul1* gene is usually embedded in class 1 integrons (also displayed by the aforementioned correlation in abundance profiles of *sul1* and *intI1* genes), whereas the *sul2* gene is usually located in small plasmids (Sköld, 2000). Thus, *sul1* and *sul2* are not necessary assembled in the same genetic cluster and can be present on different organisms independently, explaining the differences found between them. Besides, integrons are not considered mobile: they need to be inserted in other MGEs (like transposons or plasmids) to spread and replicate (Domingues et al., 2015; Ghaly et al., 2021). Plasmids usually contain genes for their self-replication, but they are also present in several copies in the bacterial cells (multi-copy plasmids) (Shintani et al., 2015). Therefore, it is more probable that genes embedded in plasmids will proliferate faster than those embedded in integrons in a nutrient-rich bacterial community. Further experiments would be needed to confirm this hypothesis.

## 4. Conclusions

Here, we showed the impact of antibiotic treatments with SMX on the abundances of marker genes coding for sulfonamide resistance (*sul1, sul2*) and class 1 integron-integrase (*intI1*) in an activated sludge, with an analytical insight from both the iDNA and exDNA pools. Collectively our results highlighted that:

- A previously developed anion-exchange chromatography method for the isolation of exDNA from wastewater environments was validated for activated sludge fractions at different biomass concentrations and volumes. Enough exDNA template (>2 μg DNA) for downstream molecular analysis was obtained with a 2-fold lower volume (0.5 L) and biomass concentration (1 g_VSS_ L^-1^), showing the robustness of the method.
- Exposure to SMX (at 1 mg L^-1^) did not impact the amount of exDNA present in the sludge. However, the exDNA concentration dropped by 70% during biomass starvation under nutrient limitations.
- The abundances of *intI1* and *sul1* genes in the exDNA fraction of the first batch exposed to 1 mg L^-1^ of SMX were positively correlated (r>0.7): most of the extracellular *sul1* genes were supposed to be embedded in the class 1 integrons.
- Starvation under nutrient limitation resulted in a decrease (of 1-2 log gene copies ng^-1^ DNA) of the *intI1* and *sul1* genes in the exDNA fraction, while *sul2* in the control flasks conversely increased (of 1 log gene copies ng^-1^ DNA).
- Under nutrient-rich conditions, the exposure to high SMX concentrations (>10 mg L^-1^) selected for the growth of microorganisms that carried the sulphonamide resistance genes *sul1* (usually located on class 1 integrons) and especially *sul2* (usually located on small plasmids) on their iDNA pool.

## Acknowledgements

This research was supported by the Spanish Government (Agencia Estatal de Investigación) through ANTARES (PID2019-110346RB-C21) project. M. Martinez-Quintela also expresses his gratitude to the same agency for awarding a research scholarship (BES-2017-080503) and for a research visit at the TU Delft for conducting this work. The TU Delft team was financially supported by the Biotechnology and Safety Program of the Ministry of Infrastructure and Water Management and the project TARGETBIO of the NWO-TTW program Biotechnology & Safety (grant no. 15812) of the Applied and Engineering Sciences (TTW) Division of the Dutch Research Council (NWO).

## Notes

### Competing Interest Statement

The authors have declared no competing interest.

