## Supplementary material for "Antibiotic resistance response to sulfamethoxazole from the intracellular and extracellular DNA fractions of activated sludge": Table S1

**Table S1.** Synthetic fragments used to obtain qPCR standard curves for 16S rRNA, *intI1*, *sul1* and *sul2* genes.

| **Gene** | **Sequence** |
| --- | --- |
| 16S rRNA | CGGCAACGAGCGCAACCCTTATCCTTTGTTGCCAGCGGTCCGGCCGGGAACTCAAAGGAGACTGCCAGTGATAAACTGGAGGAAGGTGGGGATGACGTCAAGTCATCATGGCCCTTACGACCAGGGCTACACACGTGCTACAATGG |
| *intI1* | GATCGGTCGAATGCGTGTGCTGCGCAAAAACCCAGAACCACGGCCAGGAATGCCCGGCGCGCGGATACTTCCGCTCAAGGGCGTCGGGAAGCGCAACGCCGCTGCGGCCCTCGGCCTGGTCCTTCAGCCACCATGCCCGTGCACGCGACAGCTGCTCGCGCAGGCTGGGTGCCAAGCTCTCGGGTAACATCAAGGC |
| *sul1* | CGCACCGGAAACATCGCTGCACGTGCTGTCGAACCTTCAAAAGCTGAAGTCGGCGTTGGGGCTTCCGCTATTGGTCTCGGTGTCGCGGAAATCCTTCTTGGGCGCCACCGTTGGCCTTCCTGTAAAGGATCTGGGTCCAGCGAGCCTTGCGGCGGAACTTCA |
| *sul2* | TGGAGGCCGGTATCTGGCGCCAGACGCAGCCATTGCGCAGGCGCGTAAGCTGATGGCCGAGGGGGCAGATGTGATCGACCTCGGTCCGGCATCCAGCAATCCCGACGCCGCGCCTGTTTCGTCCGACACAGAAATCGCGCGTATCGCGCCGGTGCTGGACGCGCTCAAGGCAGATGGCATTCCCG |
